# Fast and accessible morphology-free functional fluorescence imaging analysis

**DOI:** 10.1101/2025.04.15.648462

**Authors:** Alejandro Estrada Berlanga, Gabrielle Kang, Amanda Kwok, Thomas Broggini, Jennifer Lawlor, Kishore Kuchibhotla, David Kleinfeld, Gal Mishne, Adam S. Charles

## Abstract

Optical calcium imaging is a powerful tool for recording neural activity across a wide range of spatial scales, from dendrites and spines to whole-brain imaging through two-photon and widefield microscopy. Traditional methods for analyzing functional calcium imaging data rely heavily on spatial features, such as the compact shapes of somas, to extract regions of interest and their associated temporal traces. This spatial dependency can introduce biases in time trace estimation and limit the applicability of these methods across different neuronal morphologies and imaging scales. To address these limitations, the Graph Filtered Temporal Dictionary Learning (GraFT) uses a graph-based approach to identify neural components based on shared temporal activity rather than spatial proximity, enhancing generalizability across diverse datasets. Here we present significant advancements to the GraFT algorithm, including the integration of a more efficient solver for the L1 least absolute shrinkage and selection operator (LASSO) problem and the application of compressive sensing techniques to reduce computational complexity. By employing random projections to reduce data dimensionality, we achieve substantial speedups while maintaining analytical accuracy. These advancements significantly accelerate the GraFT algorithm, making it more scalable for larger and more complex datasets. Moreover, to increase accessibility, we developed a graphical user interface to facilitate running and analyzing the outputs of GraFT. Finally, we demonstrate the utility of GraFT to imaging data beyond meso-scale imaging, including vascular and axonal imaging.

## 1 Introduction

Scientific advancements in neuroscience are dependent on our capacity to observe and understand the intricate dynamics within neural tissue. Specifically, we need to analyze, and therefore discern, the activity of cells, neurites, neurotransmitters, and vasculature at both high spatial and temporal resolution, and at scale. To capture these complex dynamics across the varying scales of the brain, neuroscience has increasingly relied on advanced imaging techniques to record molecular and electrical signaling events indicative of brain function. Traditional non-invasive imaging techniques, while effective in their respective domains, often contend with limitations in terms of imaging speed, signal-to-noise ratio, and field-of-view scalability [1, 2]. Amid these challenges, two-photon excitation microscopy (2P) remains a popular choice for its unique advantage of *in vivo* deep tissue penetration with optical sectioning, enabling imaging of both structural brain dynamics and cell function [3].

Genetically encoded reporters of neural activity can reliably record changes in fluorescence that represent the underlying neural-spiking-related biophysical changes in labeled neurons. Specifically, calcium is of great interest as calcium ion concentration within the cell changes with suprathreshold (neuronal firing) activity due to the influx of calcium ions into the neuron following the depolarization associated with action potentials [4, 5]. The ability of calcium indicators coupled with multi-photon images to record *in vivo* at various scales—from dendrites and soma to widefield—at high resolutions has made calcium imaging an ideal approach for studying functional aspects of neural circuits [6, 7].

In the wake of these imaging advances, the resulting data contains ever larger fields of view at high resolution (i.e, many pixels per image) and higher temporal resolution for longer sessions (many video frames). A significant current challenge is analyzing the vast data volumes generated by such large-scale imaging efforts. Specifically, while traditional workflows relied on manual segmentation to identify single fluorescing objects and their temporal activities, manual annotation is infeasible in many modern datasets and so recent years have seen a shift towards automated analysis methods [8]. Current automated calcium imaging (CI) analysis [9–17] mainly decomposes recorded videos into two sets of variables: spatial profiles that represent neuronal occupancy, and their corresponding fluorescence time traces. These approaches, largely focused on somatic imaging, typically employ spatiotemporal regularization to reflect the sparse and well-localized nature of cell bodies. However, recent advances in CI, including finer-scale dendritic imaging and widefield imaging across the cortex, have diversified the range of observable neural structures and necessitated the development of more adaptable analysis techniques [17–19].

Relying solely on spatial structure alone can fail when the spatial morphology is complex, non-compact or does not match classical “neuron-shaped” components. In this context, the Graph Filtered Temporal Algorithm (GraFT) [17] was presented as a novel model for automated CI analysis that overcomes these shortcomings by replacing *spatial* regularization with regularization on a *graph* that captures correlations between per-pixel image time traces. GraFT is thus able to analyze a variety of tissue morphologies across many spatial scales (e.g., widefield and dendritic imaging), providing meaningful spatial and temporal components in high-dimensional imaging data. Specifically, GraFT uses the graph-based relationships between pixels to learn a set of time traces that efficiently (i.e., sparsely) represent all pixels. The result is that each learned time trace corresponds to a single spatial profile, i.e. all the pixels whose temporal activity includes this time trace. These are not constrained to be compact shapes, thus GraFT can identify components that are not spatially continuous or that span an entire FOV.

While GraFT has been successful in widefield, dendritic, somatic, and even functional ultrasound imaging (fUS) [20], the sparsity constraints and inherent matrix factorization aspect of the approach is computationally limiting for newer data taken at higher scales/resolution or at higher frame rates/longer duration. Current methods primarily deal with data limitations by downsampling [10, 11], patching [16, 17], or by simplifying the ROI detection problem (e.g., using deep learning methods) to a simple pixel-wise averaging instead of a full decomposition [21–23]. Downsampling is sub-optimal as it can reduce the ability to detect finer structures, such as axons, thinner dendrites, or spines. Patching can improve runtime by spreading the problem over more cores, but requires significant additional compute infrastructure as well as additional patch merging steps. Operating on space and time independently is also sub-optimal as summary images can be biased towards high-activity objects and averaging pixels can cause bleed-through of activity in densely labeled tissue.

In this work, we introduce a number of enhancements to the GraFT algorithm, aimed at speeding up its performance and improving its usability in the face of increasingly complex CI data. First, we integrate a more efficient solver for the L1 least absolute shrinkage and selection operator (LASSO) problem, optimizing the algorithm’s ability to accurately calculate spatial coefficients more rapidly and provide insights into parallelization parameters. Second, by incorporating principles of random projections from compressive sensing [24–26], we reduce the computationally-relevant data dimensions through a compression operation, while preserving the data quality, effectively reducing the computational workload without changing the output integrity. This approach significantly reduces the dimensionality of optimization problems within GraFT, including the calculation of weighted LASSO spatial coefficients and in the updating of the temporal dictionary. Furthermore, we implemented a Graphical User Interface that wraps the GraFT software to ensure it is accessible and easy to use, with both preprocessing capabilities and post-hoc visualization and labeling of results. We demonstrate GraFT on generate data as well as real-world cellular, vascular and axonal imaging, demonstrating GraFT’s broad applicability to neuronal morphologies. These improvements not only bolster the algorithm’s processing speed but also extend its applicability to larger and more diverse datasets, ensuring that GraFT remains a valuable alternative to other automated CI analysis tools in neuroscience.

## 2 GraFT Algorithm

We first briefly review the GraFT framework for functional imaging analysis [17]. In Sec. 3-4 we present algorithmic speed-ups to this approach.

### 2.1 Overview

Given an *N*_*x*_ *× N*_*y*_ FOV (containing *N* = *N*_*x*_*N*_*y*_ pixels overall) imaged for *T* time-steps, the GraFT (Graph Filtered Temporal) algorithm is a method designed for extracting independent fluorescing objects and their temporal activity from functional microscopy videos. Specifically, GraFT leverages the principles of graph-regularized dictionary learning (DL), an unsupervised machine learning approach that decomposes signals as sparse combinations of elementary components (i.e., the dictionary), where the underlying graph structure promotes signals that are neighbors on the graph to have similar dictionary representations. Specifically for calcium imaging analysis, GraFT decomposes the pixel-by-time fluorescence data matrix (**Y** ∈ ℝ^*N ×T*^) into a dictionary of time traces (**Φ** ∈ ℝ^*T ×M*^, where each column corresponds to the time trace of one of the *M* fluorescing components. These pixels are sparsely expressed across space, i.e. each pixel is associated with only a few times traces. The spatial expression of each component is captured in the corresponding column of the matrix ***A*** ∈ ℝ^*N ×M*^, which contains the sparse spatial coefficients that reflect the neuronal structure contributing to each time trace in **Φ**.

GraFT starts by modeling the data as the product of the spatial profiles and time traces, i.e.

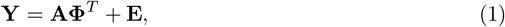

where ***E*** ∈ ℝ^*N ×T*^ represents independent sensor noise that is modeled as Gaussian. In addition to the matrix factorization likelihood, GraFT further assumes that 1) there is minimal spatial overlap between components, i.e., each pixel can contain activity from only a few of the components in the FOV, and 2) that similar pixels (measured by temporal correlations) are part of similar components. The former constraint results in a sparsity constraint over the rows of ***A***, and the latter is implemented by graph-based regularization. GraFT incorporates these priors into the overall probabilistic model by first constructing a graph over the pixels and then employing a variational expectation-maximization approach to alternate between solving for the spatial profiles ***A*** given the time traces **Φ** and vice versa. Thus, GraFT operates via three main subroutines: 1) Construction of a graph between pixels, 2) Estimation of spatial profiles given an approximation of the time traces, and 3) Estimation of the time traces given the spatial profiles.

### 2.2 Graph construction

The graph regularization is a key step in GraFT whereby the model re-organizes the data away from the spatial domain into a correlation-driven graph. GraFT accomplished this by constructing a graph such that each node represents a single pixel and each edge weight connecting two nodes (pixels) represents the similarity of the two corresponding pixels’ time traces. The graph captures the intrinsic structure of the data independent of pixel location in the FOV, allowing for the exploitation of the temporal correlations present in the imaging data without explicit morphological constraints. Thus two pixels on the same dendritic branch can be linked on the graph, even if they are spatially far apart. The graph is constructed by first computing the weighted graph affinity matrix ***W***. Each element *W*_*ij*_ between pixels *i* and *j* is computed with a Gaussian kernel as

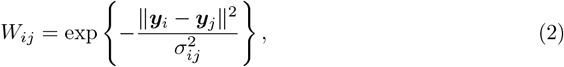

where ***y***_*i*_ is the time trace of pixel *i* and the bandwidth *σ*_*ij*_ = *σ*_*i*_*σ*_*j*_ is chosen according to a local adaptive approach [27]. The graph is constructed via a k-nearest neighbor search such that the resulting graph is sparse (each node is connected to only a small number of other nodes). This prevents many weak connections from over-powering smaller numbers of strong connections in larger FOVs with more pixels. Finally, the rows of *W* are normalized to yield the diffusion graph filter ***K*** as ***K*** = ***D***^*−*1^***W***, where ***D*** is a diagonal degree matrix with entries summing each row of ***W***.

### 2.3 Model inference

After constructing the graph kernel ***K***, GraFT then alternates between updating the spatial profiles ***A*** and the time traces **Φ**. The update over ***A*** is accomplished via a regularized least-squares minimization based on sparse coding, where the sparse codes are aligned to the graph. This optimization is performed through the Re-Weighed *ℓ*_1_ Graph Filtering (RWL1-GF) optimization procedure. In RWL1-GF, the component time traces (columns of **Φ**) that best represent each pixel are inferred simultaneously across all pixels while promoting minimal overlap through a non-negative weighted LASSO. The LASSO weights, which promote or suppress a given component from appearing at a given pixel, are then updated at each iteration based on how many of the pixels that are neighbors on the graph are also well represented by that component.

Mathematically, RWL1-GF iterates between the two steps of estimating the sparse spatial coefficients and updating the weights

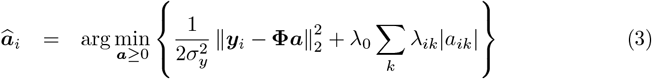

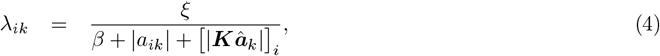

where ***a***_*i*_ is the strength of all spatial profiles at pixel *i*, ***λ***_*i*_ are the LASSO weights for the coefficients, *β* is a weight parameters, and *λ*_0_ is included for numerical stability.

The weights ***λ*** in RWL1-GF incorporate non-local spatial information into per-pixel solutions and maintain sparsity via applying the graph-based filter to the current estimate of the spatial coefficients 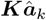 [15]. At a high level, if the spatial profile coefficients for component *k* of the *i*^*th*^ pixel’s graph neighbors are high, the corresponding weight *λ*_*ik*_ will be low, encouraging that component to also be active in pixel *i*. Conversely, if the neighbor’s spatial profile coefficients have low weights, the LASSO weight for that component at pixel *i* will be high and bias that component away from being active at that pixel.

Once the spatial profiles ***A*** are updated based on the time trace dictionary, the dictionary **Φ** is then updated based on the current estimate of ***A***. The complete dictionary update for the estimate at the *l*^*th*^ update iteration 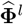 is given by

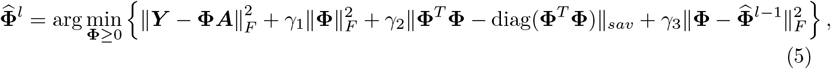

where here the SAV (sum-absolute-value) norm is defined as 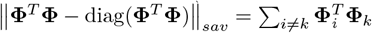. The first term in this objective is a data fidelity term such that the dictionary and the sparse coefficients faithfully reconstruct the data. The second term regularizes the magnitude of the time traces. The third term is a regularization which aims to prevent distinct components having highly correlated time traces. The final term is an optimization smoothness term preventing subsequent algorithmic iterations from changing too drastically. The parameters *γ*_1_, *γ*_2_, *γ*_3_ trade-off between the relative importance of each term to the optimization solution. Put together, the full GraFT algorithm is summarized in Algorithm 1.

## 3 Practical LASSO Optimization

### 3.1 Problem formulation

A key step in GraFT is the constrained, weighted LASSO, a regression technique that imposes a non-uniform *ℓ*_1_ norm over the regression coefficients. This constraint encourages sparsity in the model by effectively shrinking most coefficients to zero, thereby performing simultaneous variable selection and estimation. LASSO minimizes the residual sum of squares subject to the sum of the absolute values of the coefficients being less than a fixed value. The standard LASSO optimization problem can be expressed as:

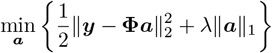

#### Algorithm 1

GraFT DL

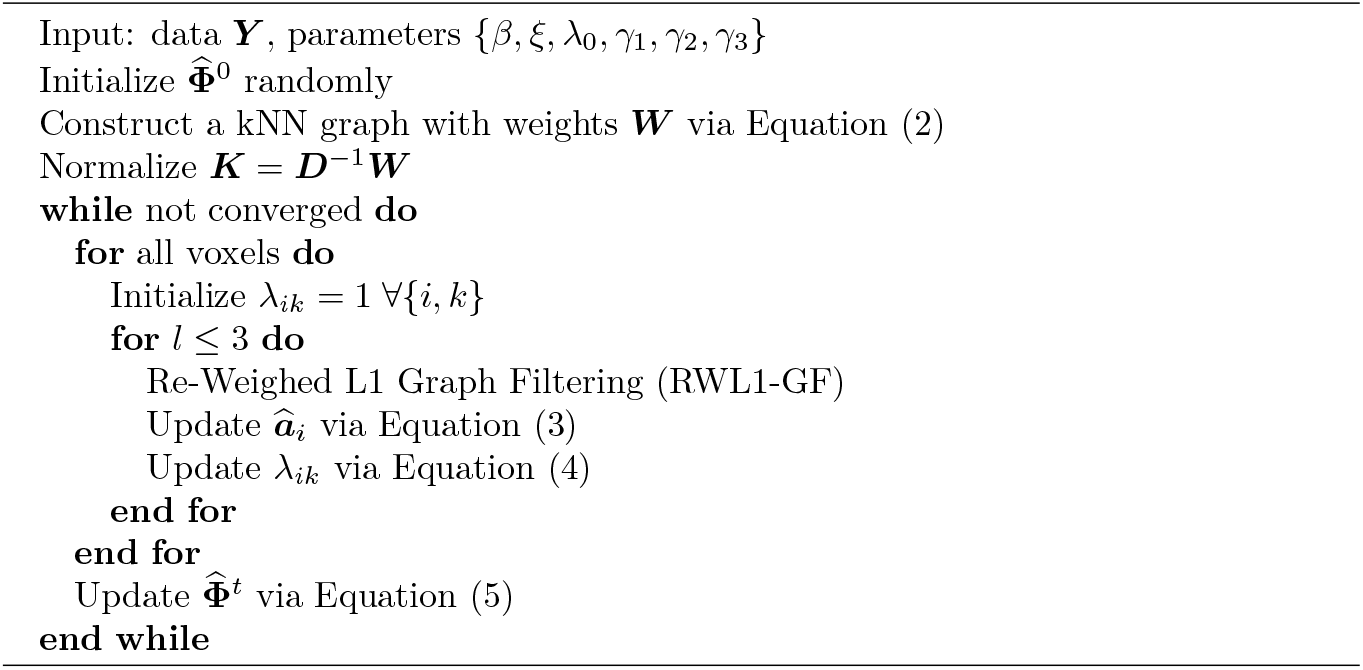

where ***y*** ∈ ℝ^*n*^ is the response vector (i.e. the pixel’s time trace), **Φ** ∈ ℝ^*n×p*^ is the matrix of predictors (i.e. the time trace dictionary), ***a*** ∈ ℝ^*p*^ is the vector of coefficients (i.e. the pixel’s “expression” of each of the traces in the dictionary), and *λ* ≥ 0 is a regularization parameter. Here, ∥***a***∥_1_ is the *ℓ*_1_ norm of the coefficients, defined as 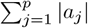. The *ℓ*_1_ norm can be expressed as a linear combination of auxiliary variables, which allows the problem to be formulated as a quadratic program (QP) problem with linear constraints.

This reformulation not only simplifies the LASSO optimization into a standard form but also allows leveraging sophisticated, well-studied QP solvers that can handle large problem instances efficiently. Moreover, such solvers provide robust performance in terms of numerical stability and convergence guarantees, which is crucial as neural imaging datasets grow in size and complexity. To formulate this as a quadratic programming (QP) problem, we introduce auxiliary variables *u* ∈ ℝ^*p*^ such that:

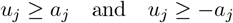

for *j* = 1, …, *p*. This reformulates the *ℓ*_1_ norm as a linear combination, allowing us to express the problem as a QP problem with linear constraints:

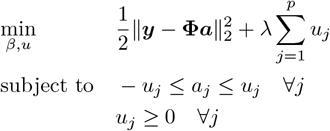

*u*_*j*_ ≥ 0 ∀*j*

Rewriting the problem in standard QP form:

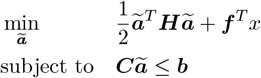

where 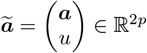, *H* is a block matrix derived from **Φ**^*T*^ **Φ**, *f* is a concatenation of *−* **Φ**^*T*^ ***y*** and *λ***1**, where **1** is a vector of all ones, ***C*** captures the linear constraints and ***b*** represents the bounds ensuring non-negativity and other problem-specific requirements.

The formulation of the LASSO problem in the framework of QP is crucial as it enables fast, reliable, numerically robust solvers to be used. It enables the use of state-of-the-art solvers that are optimized for large-scale problems, taking advantage of well-established, highly optimized QP solver libraries.

### 3.2 LASSO and QP Solvers

Choosing the appropriate QP solver for the given LASSO is crucial for the optimization and utility of the GraFT algorithm. The performance of QP solvers can significantly vary depending on the size and sparsity of the constraint matrix, making it essential to select the right approach for the specific problem at hand. Two principal iterative methods include *active-set* and *interior-point* solvers. Originating from the simplex method for linear programming [28], active-set methods (ASM) iteratively select and adapt a set of constraints that directly impact the solution in a given iteration. These methods are particularly beneficial in applications that require warm-start strategies, such as sequential quadratic programming (SQP) and model predictive control (MPC). Some drawbacks to ASM include the potential exponential growth in worst-case complexity with the number of constraints and issues like active-set cycling when constraint qualifications are not met [29].

The Interior Point Method (IPM) originally gained popularity as a polynomial-time algorithm for linear programming (LP) and was later extended to general convex optimization problems. Unlike ASM, IPM models iterate within the feasible region of the problem, approaching the solution from the ‘central path’ [30]. In the primal case, IPM methods use a *barrier function* to ensure that iterations remain within the feasible region and gradually reduce this barrier to zero as the solution is approached.

A barrier function for the constraints ***Ca*** ≤ ***b*** can be written as follows:

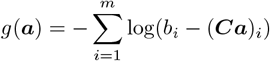

where *m* is the total number of constraints. The modified objective function becomes:

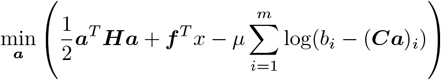

where *µ* is the barrier parameter, which is systematically decreased between iterations, smoothly guiding the solution towards the boundary of the feasible region and ultimately to the optimal point. This approach allows for significant progress per iteration, though each iteration is computationally expensive.

The time complexity of IPMs is typically *O*(*n*^3^) per iteration of input size *n* due to the matrix factorization involved in each step. However, because IPMs require fewer iterations to converge than ASM, they are efficient for large-scale problems. On the other hand, the time complexity of ASM is typically *O*(*kn*^2^), where *k* is the number of iterations. Each iteration involves solving a linear system of equations, which is computationally efficient for small to medium-sized problems but may struggle to scale.

The choice between ASM and IPM depends on the specific characteristics of the problem, including size and sparsity of the constraint matrix. An MPC study comparing these methods found that ASM is computationally faster with fewer variables and constraints, whereas IPM scaled better with larger, more complex problems [31, 32].

Conversely, ASM is suitable for smaller problems and adjusts its working set of constraints iteratively to find the optimal set, which can be computationally intensive for larger datasets. The ability to selectively choose between IPM and ASM solvers can reduce the computational burden, thereby leading to faster convergence.

Thus, the selection of a QP solver has a profound impact on the efficiency and performance of GraFT algorithm. ASMs are advantageous for problems requiring warm-start strategies and small to medium-sized datasets (for example when GraFT is parallelized over patches of the FOV rather than the full FOV), whereas IPMs are better suited for large-scale problems due to their fewer iterations and robust performance. If the GraFT pipeline is intended to run repeatedly on growing datasets, the warm-start strategies available to ASM may reduce computation times. In contrast, for very large datasets characteristic of wide-field imaging or whole brain recordings, IPM’s better scaling properties are likely to outweigh the per-iteration cost. Adjusting the solution choice based on the size, complexity and required runtime efficiency of the data significantly enhances the overall performance of the algorithm. Here, we have implemented both solvers for the GraFT solution and compare the solvers in Sec. 6.

## 4 Compressive GraFT for fast processing

While judicious selection of LASSO optimizers enables a computation speed-up, further speedups can be obtained by solving an inherent bottleneck given by the size of the data. Projections of the data onto spatial and temporal components (i.e., inner products along the space or time dimension) must come in multiples of the number of pixels *N* or time-points *T*. To further speed up GraFT we thus turn to computation in a compressed representation. Specifically, GraFT operates on the assumption that each pixel’s time trace is sparse in the dictionary of all the time traces of the components in the video (i.e., minimal spatial overlap). Sparse signals can be highly compressed into a very low-dimensional subspace without extensive distortion of their geometry. This property has been extensively studied in the compressive sensing literature and can be quantified as follows: If ***y***_1_ and ***y***_2_ are both *K*-sparse in a representational dictionary **Φ**, then a linear projection **Ψ** that does not distort distances satisfies, for all *K*-sparse ***y***_1_ and ***y***_2_,

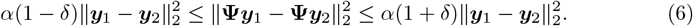

This condition is known as the Restricted Isometry Property (RIP) and has parameters *α*, which measures the inherent scaling of the operator **Ψ** and *δ*, a value between zero and one that captures how distorted the low dimensional representation can be. For *δ* = 1 two distinct points can map onto each other, while *δ* = 0 represents a perfect preservation of geometry. The RIP ensures that relative distances between sparse points are not distorted, and can hold for large compression rations in the projection **Ψ**. The preserved geometry means that many computations can be performed in the compressed space, saving computation time by operating over smaller vectors.

We implement random projections in the GraFT pipeline by compressing the data along the temporal dimension: 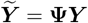. The projection matrix 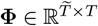 reduces the full time trace from length *T* to length 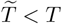. In the context of our implementation, both **Φ** and corresponding temporally-compressed input 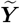 are computed once, prior to the initiation of the main dictionary learning loop. The core two steps that GraFT alternates between are then minimally changed from a mechanical standpoint, operating over the compressed version of the data instead of the full version. The first primary savings comes in the form of a reduced space in which to estimate the coefficients at each LASSO step:

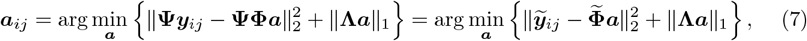

where 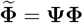is a compressed version of the component time traces. Note that since ***a***_*ij*_ is sparse, we expect that the compressed data will be sufficient to recover ***a***_*ij*_. The second savings comes in the update for the compressed dictionary 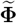

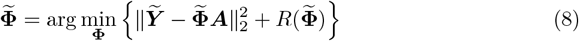

where *R*(**Φ**) represents the additional regularizers in Equation 5. This problem is over a much lower dimensional space (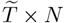 instead of *T × N*), greatly speeding up the computation. This compressed approach results in the estimate of ***A*** and 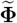, i.e., we obtain the spatial profiles as before but only a compressed version of the time traces. To recover the full time traces, we can solve Equation 5 once at the end of the optimization using the inferred ***A*** values and the uncompressed data ***Y***.

## 5 GraFT Graphical User Interface

To enhance the usability and accessibility of the GraFT algorithm for the community, we developed a dedicated graphical user interface (GUI) that consolidates preprocessing, parameter setup, algorithm execution, and results visualization into a single workflow (Fig. 1). The GUI is offered in two forms: a stand-alone compiled application for Windows systems and a MATLAB-based app compatible with MATLAB R2024a or newer^1^. Both versions are designed to simplify adoption of GraFT for imaging analysis across modalities and spatial scales.

**Fig 1.**
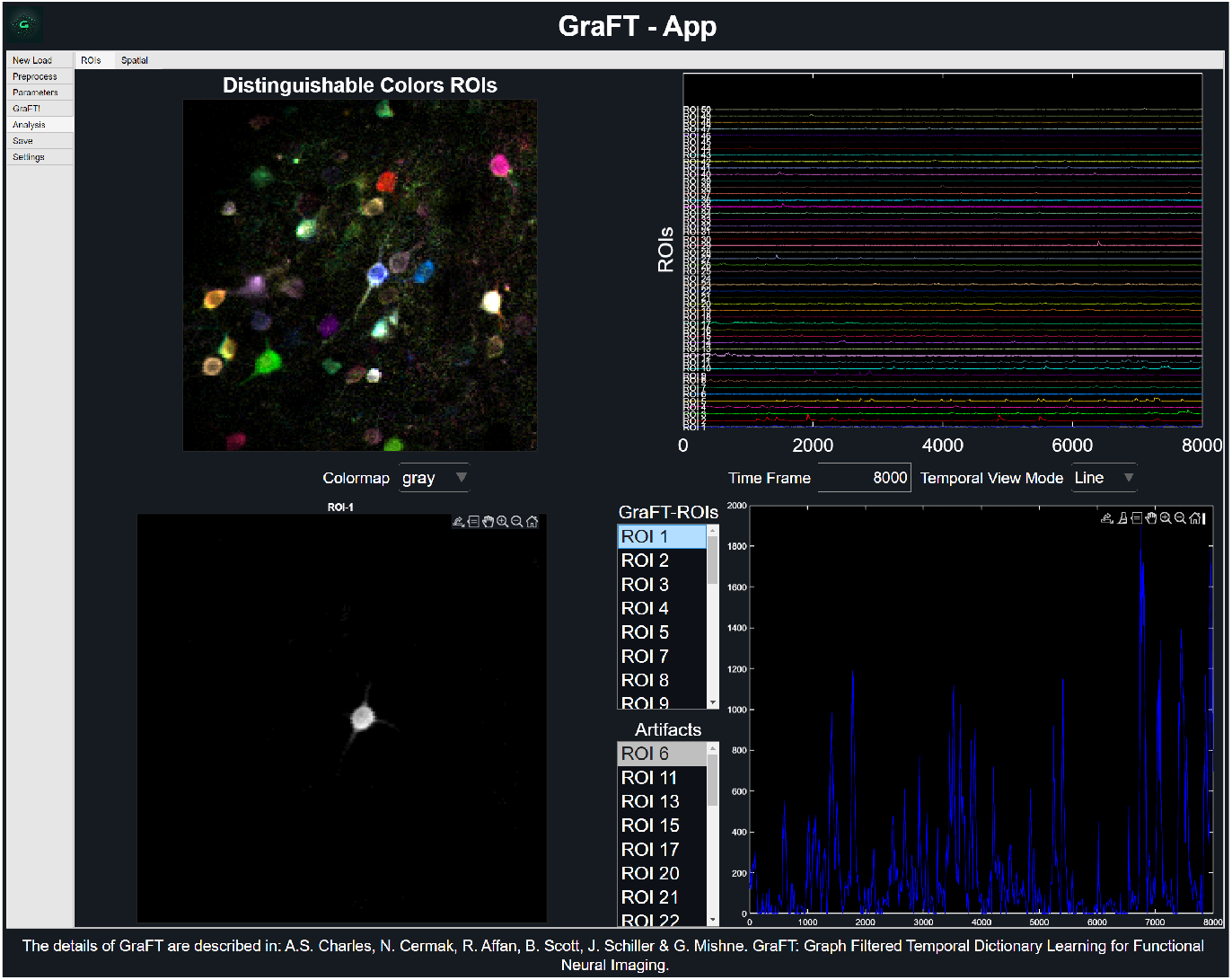
Results display from GraFT-App. (A) The distinguishable colors ROIs figure shows the overlay of all of the dictionary elements found, with their color matching their time trace in (B).

Upon launching the GUI, users can load datasets in multiple file formats, including *.h5, *.mat, as well as common image formats (*.tiff). The interface supports essential preprocessing steps such as interactive cropping for visually selecting regions of interest, manual or semi-automated mask selection to define or exclude specific areas (e.g., in sparse dendritic imaging), motion correction to address drift artifacts, and wavelet-based denoising to improve signal clarity.

Prior to running the GraFT algorithm, users can save the algorithm parameter file with project name and author identifiers for reproducibility. Previously saved parameter configurations can also be loaded or shared with other users separately from the project, which is ideal for facilitating reproducible analyses and working collaboratively. After running GraFT algorithm, the analysis tool within GraFT-App allows for quick viewing of all identified spatial profiles and time traces, selected individual or groups of spatial profiles along with their time trace. We also include labeling functionality between real neural components and artifacts.

## 6 Experimental results

In this section, we evaluate GraFT’s computational efficiency and reconstruction accuracy. We compare two LASSO solvers—quadprog (IPM) and MPC (ASM)—using the NeuroFinder dataset, showing that although both yield similar outputs, quadprog’s runtime increases nearly twice as fast with added dictionary elements. We also assess the effect of compression (2×–1024×) on time trace and spatial profile reconstruction, demonstrating substantial speedups and reduced RAM usage with minimal loss in accuracy. Finally, we demonstrate GraFT on two non-cellular morphologies and modalities: widefiled imaging of wide-field pial arterioles and two-photon axonal imaging.

### 6.1 QP Computational Results

First we isolate and test the differences in using different solvers for the LASSO optimization. Recognizing the limitations of each solver in certain scenarios, we compared the default GraFT IPM solver quadprog and its ASM counterpart. For this, we transitioned to a specialized solver, the mpcActiveSetSolver from MATLAB’s MPC Toolbox [33], which employs the KWIK algorithm [34] for efficient active-set adjustments. This solver optimizes the selection process of constraints likely active at the optimal solution, thereby expediting convergence and reducing computational load [35]. The NeuroFinder dataset [36] was motion corrected using the Non-Rigid Motion Correction algorithm (NoRMCorre) and denoised using simple time trace wavelet denoising [37]. The processed (512 × 512) pixels *×*8000 time-frames NeuroFinder data was then cropped into various sizes for testing.

We tested the GraFT algorithm runtime on (210 × 210, 160 × 160, 80 × 80, 65 *×* 65) pixel × 8000 time frames with IPM and ASM solvers. In all data size comparisons, the ASM solver with MPC was faster than the IPM solver with little to no differences in quality of temporal and spatial results. Each data size was run 12 times, and runtime was tracked for each. The relative difference between the two solvers grows with decreasing data size (Fig. 2A).

**Fig 2.**
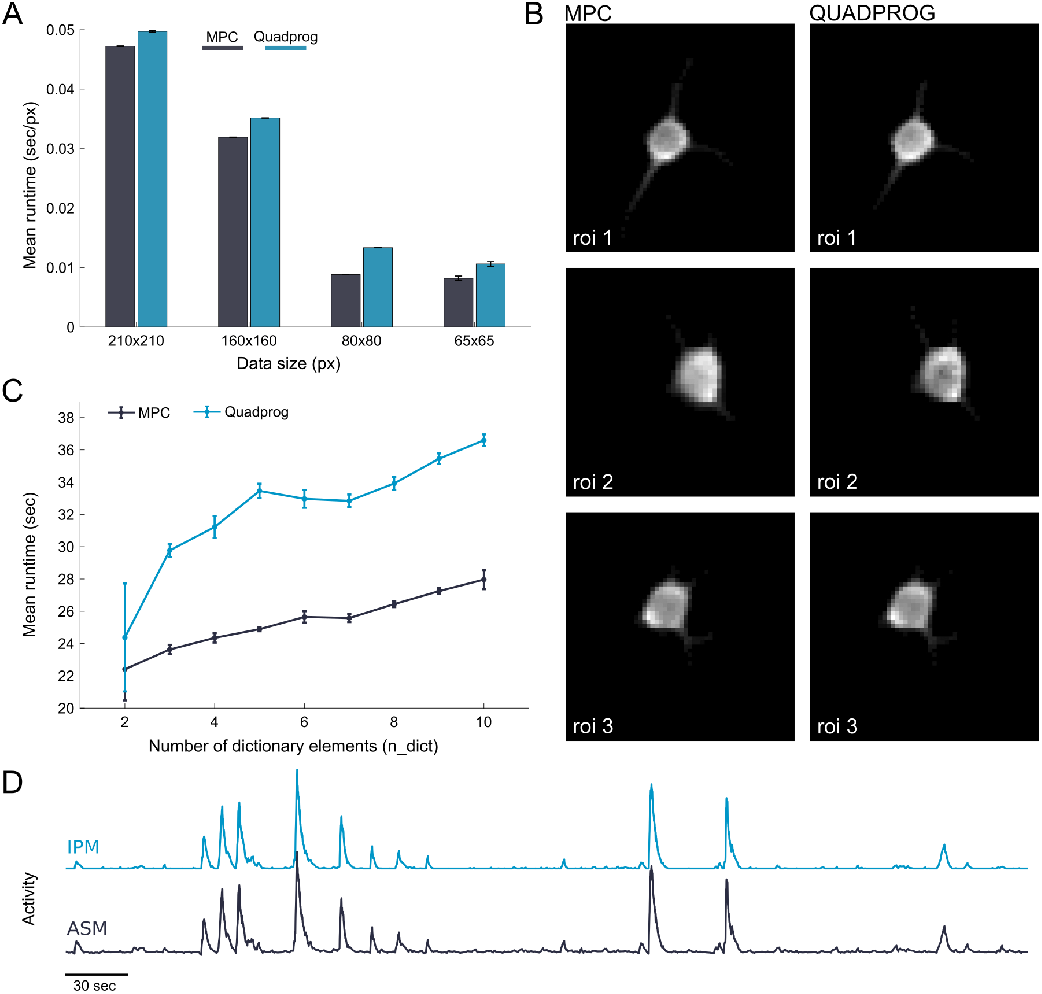
Comparison of ASM (MPC) and IPM (quadprog) solvers. (A) Runtime of solvers normalized by window size (pixel area). (B) Sample ROIs spatial results of the NeuroFinder dataset between both solvers. (C) Comparison of runtime between MPC and quadprog solvers by varying dictionary elements over 50×50 pixel area using patchGraFT. (D) Sample time trace of the same ROI (#3) from IPM and ASM solvers.

We further compared runtime differences of ASM and IPM solvers by varying the number of dictionary elements over a consistent (50 × 50) pixel window. In our patchGraFT implementation, as the number of dictionary elements increased, the runtime for the quadprog solver rose more steeply than that of MPC (Fig. 2B). Linear regression on the experimental data revealed that the MPC solver’s runtime increased by approximately 0.63 seconds per additional dictionary element, whereas the quadprog solver’s runtime increased by about 1.18 seconds per additional dictionary element.

This indicates that for every extra dictionary element (i.e. additional neural component to be identified), quadprog incurs nearly twice the additional computation time of MPC, highlighting a significant advantage for warm-start strategies with small datasets via ASMs solvers.

### 6.2 Comparison between compressed an uncompressed GraFT

We evaluated the computational and accuracy performance of compressed GraFT using a 490 × 490 × 100 *µ*m × 9,000 time points (equivalent to 3 minutes) generated with NAOMi’s default parameters (details in App.A). The degree of compression varied from 2× to 1024 ×, and we used both normalized and non-normalized sketching matrices. First, we examined the preservation of time trace reconstruction in compressed GraFT using a non-normalized sketching matrix (*λ* = 0.7), by leveraging the known ground truth events present in the synthetic NAOMi dataset. For each trace, we calculated the correlation coefficient between output GraFT trace and input NAOMi ground truth. Each GraFT trace was tagged by its highest correlation coefficient, allowing multiple GraFT traces to map to the same NAOMi trace. Compiling the trace correlation tags into a histogram (Fig. 3A) displays in skewed right behavior. As compression level increases up to 16 ×, the trace correlation histogram begins to flatten at lower correlation levels, and GraFT identifies more traces. However, as compression level further increases up to 512 ×, increasingly many traces, previously unidentified traces with low correlation (*ρ <* 0.3) to the ground truth NAOMi data are found, leading to an increased amount of raw identified traces.

**Fig 3.**
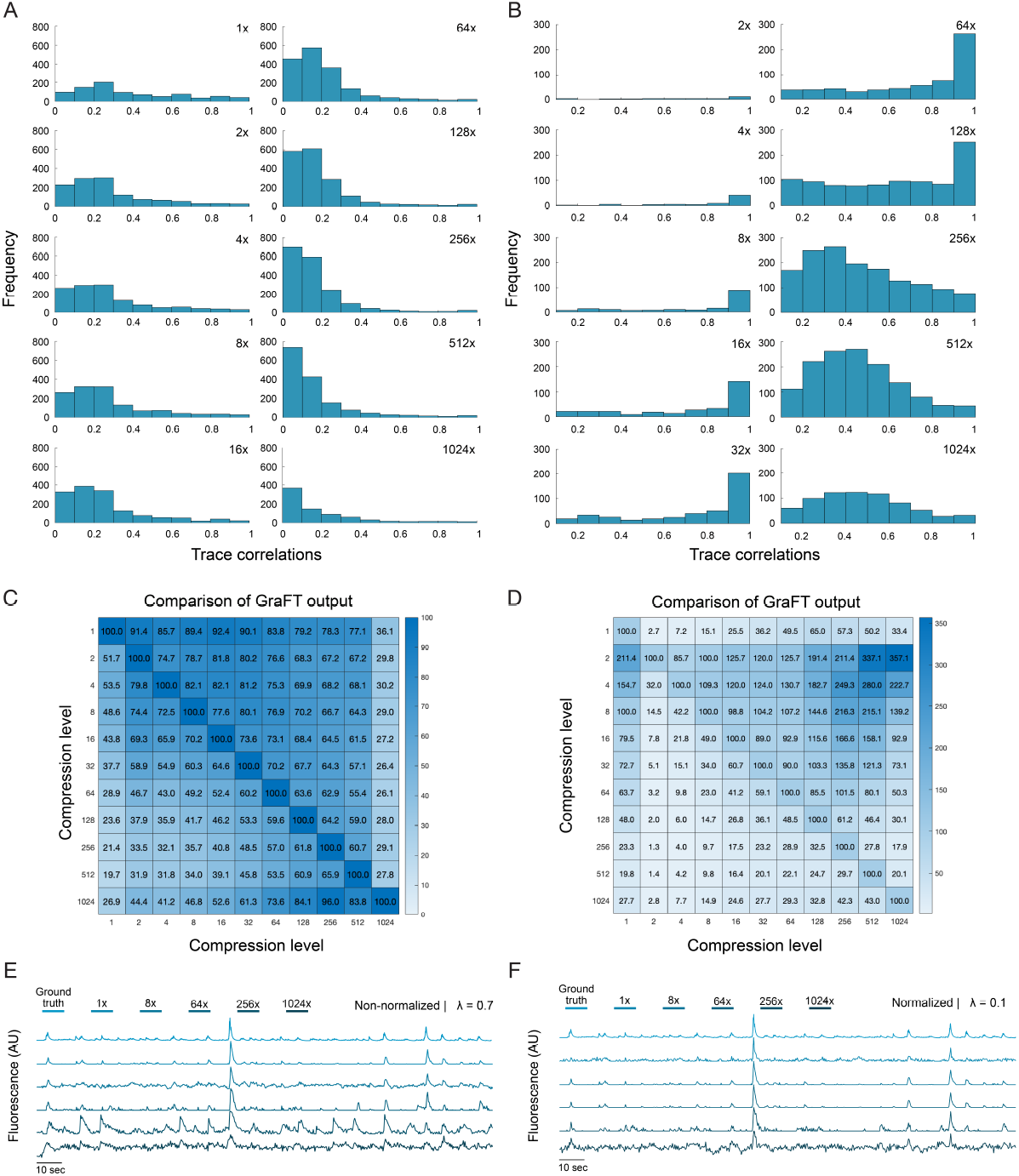
Effects of time compression on temporal components identified by non-normalized and normalized GraFT. Histogram of highest time trace correlations to the NAOMi ground truth for (A) non-normalized GraFT and (B) normalized GraFT. Heatmap for (C) non-normalized and (D) normalized GraFT results across compressions levels. Each entry is the percentage of traces in the row-wise compression result that correlated at *ρ >* 0.5 with the column-wise result. Example highly correlated traces (*ρ >* 0.7 for all compression levels) in (E) non-normalized GraFT and (F) normalized GraFT

Examining individual time trace profiles, we see preservation in trace correlations across compression levels. Strong trace correlations (Fig. 3E) retain high correlation across increasing compression magnitudes. Moderate trace correlations to the ground truth trace gradually decay in correlation, with the most noticeable drop in correlation starting at 64 × compression (Fig. S1A).

The output traces from compressed GraFT were also compared against other GraFT compression level outputs to assess self-agreement of the GraFT algorithm (Fig. 3B).

Specifically, we select one GraFT output as the pseudo-ground truth (listed on the y-axis) and another GraFT output as the baseline run (listed on the x-axis). The correlations between this pseudo-ground truth and comparison were calculated per trace.

We performed the same highest correlation tagging procedure as before, but computed the number of GraFT traces with a correlation *ρ >* 0.5 in the comparison as a percentage with respect to the number of traces identified in the y-listed compression level GraFT run. With this method, *>* 100% relationships indicate that more traces are found in the comparison than the ground truth and that multiple traces may map to the same ground truth trace. Higher percentages in the heatmap correspond to higher levels of self-agreement in the output GraFT dictionary, and we see consistency across compression levels. This suggests that increasing the compression level identifies similar traces as lower compression levels. Using the first row, we see that at least 75% of time traces identified by uncompressed GraFT are also identified with correlation coefficient *ρ >* 0.5 in non-normalized, compressed GraFT up to 512 × compression. Additionally for all combinations of levels up to 64 × compression, lower compression levels have at least 70% agreement with higher compression levels (upper triangular). This indicates that higher compression levels consistently identify similar traces to lower compression level but, when combined with the observation that the lower triangular has lower agreement percentages, also indicates that a higher raw number of traces are identified.

For comparison of normalized GraFT time trace reconstruction, the sparsity parameter was changed to account for compression (from *λ* = 0.7 to *λ* = 0.1). With the lower *λ* parameter, notably fewer traces are identified overall (Fig. S1C), but the remaining traces identified by normalized GraFT have higher correlation to the ground truth across compression levels (Fig. 3B). However, we see that the percentage of highly correlated traces in respect to uncompressed GraFT as the pseudo-ground truth has a maximal performance around 65% at 128 × compression (Row 1, Fig 3D). Although the maximal performance is less than with non-normalized GraFT (which had maximal performance around 80% at the same 128 × compression level), the strength of the correlation is much higher for the identified traces (Fig. 3B). The individual time traces demonstrate similar preservation of trace correlation strengths across compression levels, with more stable preservation of mid-correlated traces (Fig. S1B).

### 6.3 Comparison of Spatial Profile Reconstruction in GraFT

For evaluation of spatial profiles, we used a 500 × 500 *×* 100 *µ*m × 20,000 time point at 30 Hz (approximately equivalent to 11 minutes) synthetic NAOMi dataset for analysis. The individual spatial profiles generated from non-normalized GraFT preserve important structures but have increased background noise as compression level increases. We note that time traces with *ρ <* 0.5 correlation to the ground truth still have relevant spatial information at 64 × compression (Fig. 4A). However, the when examining the full volume, there are strong patch boundaries (where individual regions of the image are brighter or darker than neighboring patches) and a reduction in captured cells (Fig. 4B). Spatial profile visualized can be improved via post-processing techniques adjusting filter sensitivity or contrast, indicating that the “missing” spatial profiles are still identified but experiencing pixel intensity modulation from the introduction of the sketching matrix (Fig. 4C).

**Fig 4.**
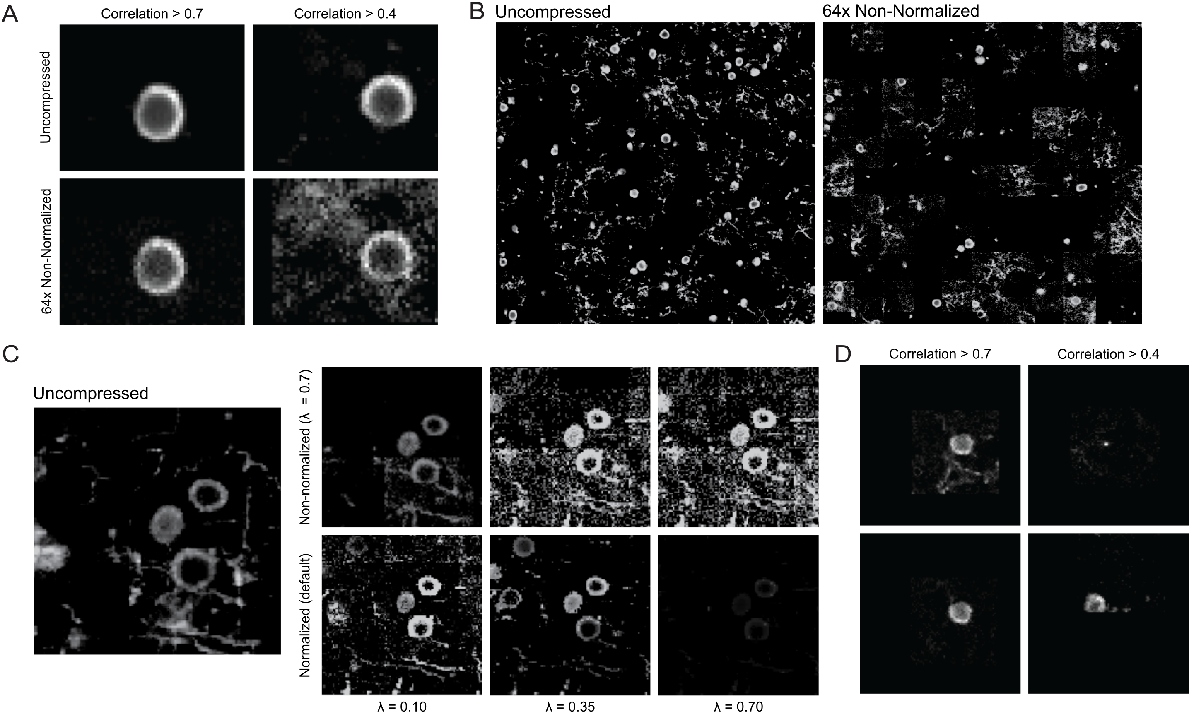
Effects of time compression on spatial profiles identified by non-normalized and normalized GraFT. (A) Plot of spatial profiles comparing uncompressed GraFT and 64 ×, non-normalized GraFT which had highly correlated time traces (*ρ >* 0.7) and medium correlated time traces to the ground truth (0.4 *< ρ <* 0.7). (B) Plot of all spatial profiles with corresponding time trace correlations *ρ >* 0.5.(C) Spatial profile comparison within a defined ROI between non-normalized, *λ* = 0.7, 64 × compressed GraFT and normalized, 64*×* GraFT. The non-normalized ROI was post-proceesed using different filtering parameters to improve visualization, while the normalized ROI used the default visualization parameters across different sparsity *λ* coefficients. (D) Comparison of non-normalized, *λ* = 0.7, GraFT (top) and normalized GraFT, *λ* = 0.1, GraFT (bottom) spatial profiles comparing to the time traces shown in Fig. 3

**Fig 5.**
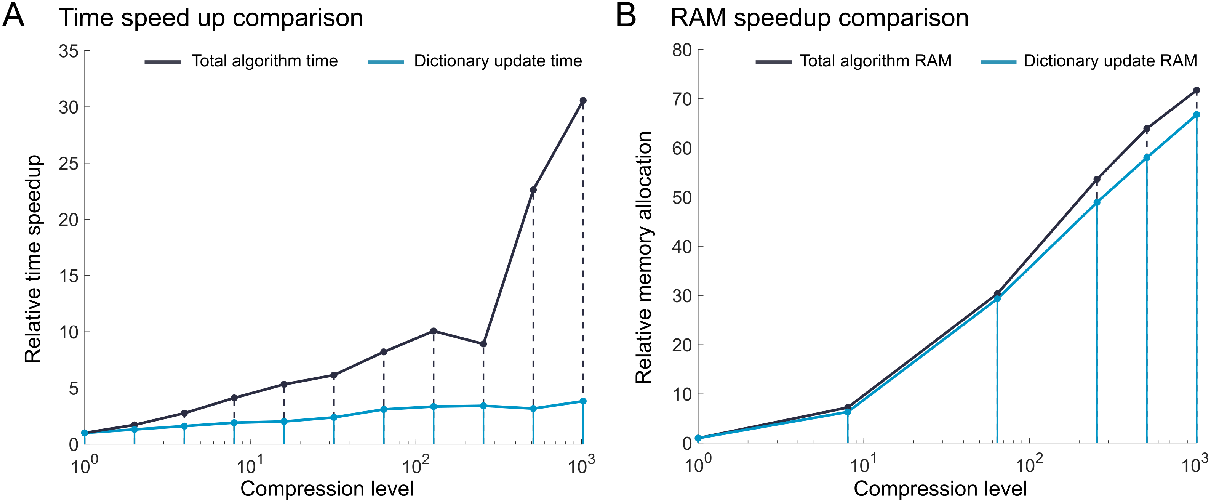
Time speedup and RAM allocation improvement by compression level. (A) Relative time speedup as a function of compression. (B) Relative RAM usage as a function of compression

By implementing normalization among the sketching matrix, we are able to see less noisy spatial profile visualization with the default display parameters. Additionally, we are able to directly visualize the effect of modifying of the sparsity coefficient, *λ*, where decreasing the coefficient identifies more spatial profiles, and correspondingly use the results from *λ* = 0.1 for subsequent analysis on spatial profiles corresponding to the time traces highlighted in Fig. 3. We see that non-normalized GraFT results have more background contamination in the case of the higher correlated trace and worse recognition of a cell ROI for the mid correlated trace (Fig. 4D).

### 6.4 GraFT Speedup

We used the inbuilt MATLAB profiler to evaluate the run time and RAM usage of GraFT at different compression levels. In addition to extracting the metrics for the total algorithm, we also extracted the metrics of the dictionary update step (which was expected to show the most improvement). The relative metric at each compression level was calculated by dividing the compressed metric by the uncompressed metric. We see a comparable relationship between relative algorithm speedup and compression level across the dictionary update step (8 × relative speed up at 64 × compression). Generally, relative speed up followings a root based trend, where speed up is equivalent to the square root of compression level at the time intensive step. The notable speed up in terms of total algorithm runtime is less significant, stabilizing at 3 × relative speed up at 64× compression. On the other hand, relative RAM usage demonstrates consistent root based trends in both the time intensive and overall algorithm step.

### 6.5 Analysis of diverse morphologies with GraFT

GraFT is designed as a morphology-agnostic approach for functional imaging analysis. Here we highlight two non-somatic imaging datasets, to complement the cellular data in previous sections.

#### Vasculature widefield data

We use GraFT to analyze calcium imaging of pial arterioles of a mouse at rest from the study in [38], in order to identify sub-structures within the pial arteriole network and their corresponding time traces. Using a masking option, we restrict the analysis only to pixels that fall within the mask of the pial arterioles, resulting in 6197 pixels by 2223 time frames (see supplementary for details on mask construction). In Fig. 6A we present the extracted spatial profiles. GraFT extracts five profiles that all exhibit bilateral symmetry and partition the arterial network into distinct regions. Note that due to the graph-based nature of GraFT, a spatial profile need not be contiguous and can be composed of multiple segments. In Fig. 6B we present the time trace dictionary, where each trace corresponds to one of the extracted spatial profiles in Fig. 6A. Finally, Fig. 6C shows Pearson’s correlation across the spatial and temporal components, exhibiting low correlation between the spatial profiles. Profiles that are closer together spatially are more correlated temporally.

**Fig 6.**
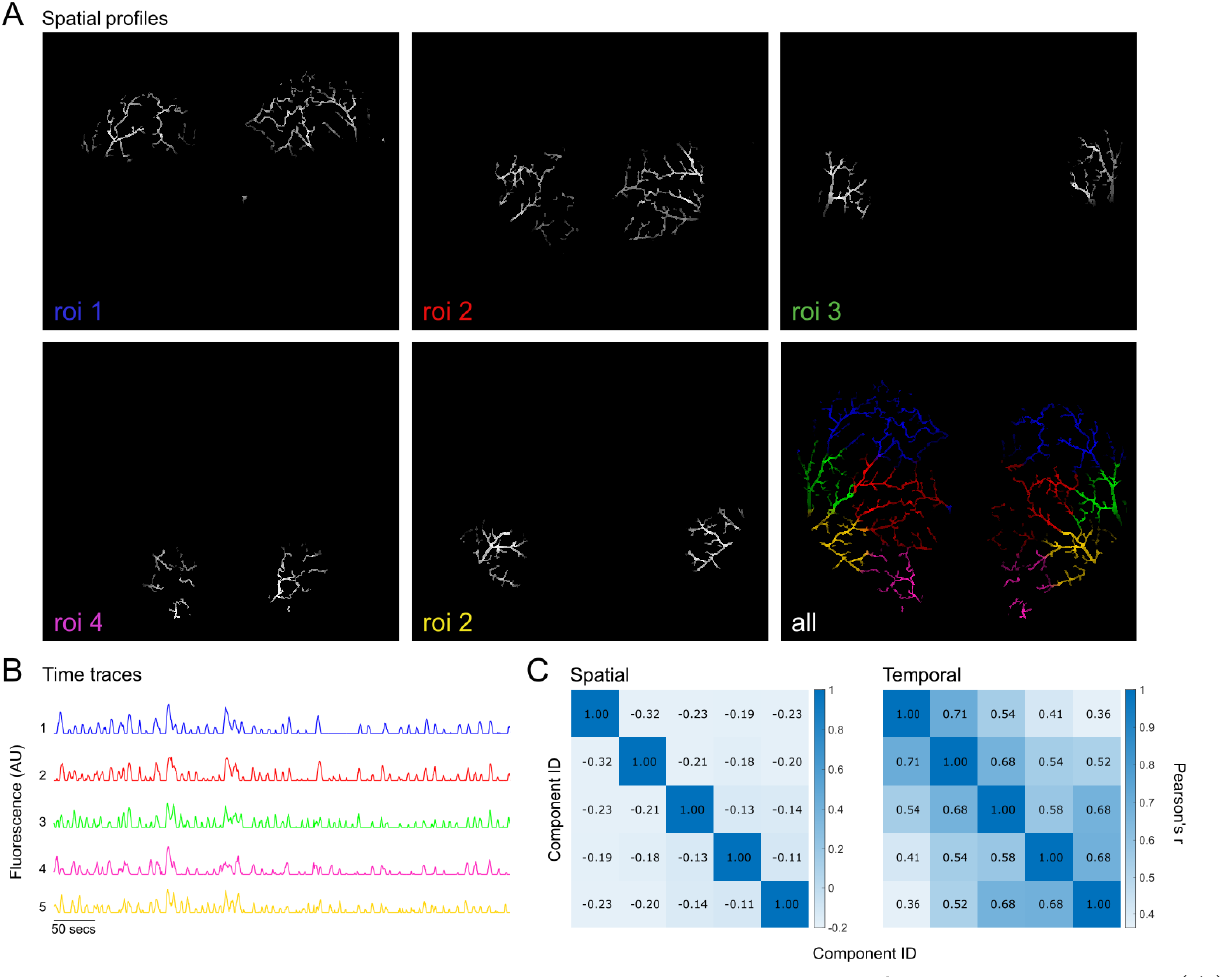
GraFT algorithm results on calcium imaging of pial arterioles. (A) Spatial profiles found along with an overlay of all five ROIs with distinguishable colors. (B) Respective time traces of the spatial profiles detected. (C) Component spatial and temporal linear correlations using Pearson’s correlation coefficient (r).

#### Axonal data

We use GraFT to analyze calcium imaging of cholinergic basal forebrain (CBF) axons imaged in the Primary Auditory Cortex (A1) of the mouse. Data was collected as in [39], resulting in a 312 × 192 *µm*^2^ area at a frame rate of 15.63 Hz. The mean image of a region of the field of view as well as the overlay of the primary components identified by GraFT are shown in Fig. 7A (left and right, respectively). GraFT extracts 4 spatial profiles that are distributed across the field of view (Fig. 7B), reflecting the distributed nature of axonal trees in cortical target regions. For the same field of view, suite2p [11] ROI segmentation is an order of magnitude larger (48 components total) which would require significant post-processing to merge segments belonging to the same axonal track. The fluorescent trace of each spatial GraFT component is displayed in Fig. 7C for the example imaging session. Large increases in fluorescence reflect ongoing cholinergic tonic activity (highlighted with the average activity of the component and its corresponding mean image). Tonic activity is component-specific indicating that GraFT, identified distributed spatial profile with distinct temporal profiles.

**Fig 7.**
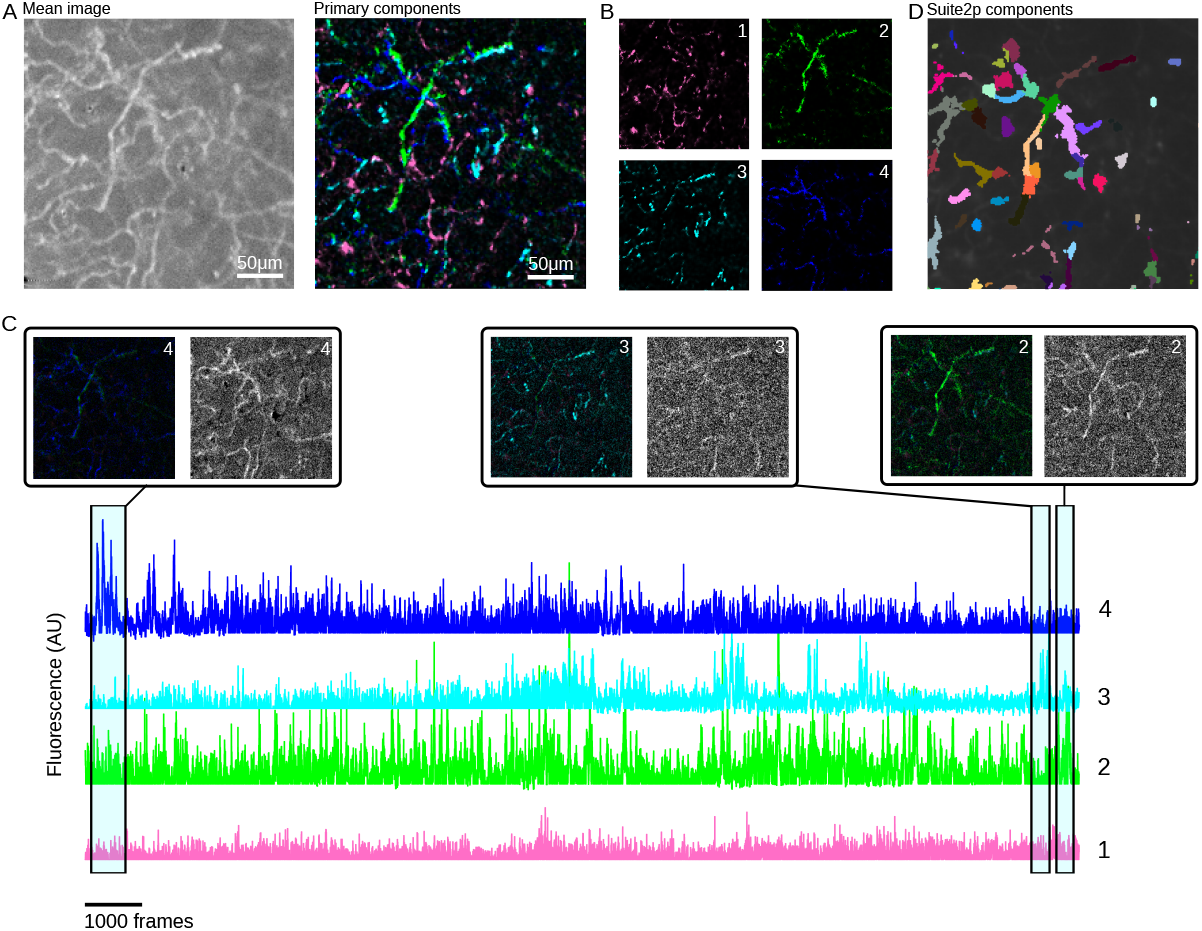
Results of GraFT on axonal imaging. (A) Mean image (left) and four identified components (right) show that GraFT finds networks of axons across the full field-of-view. (B) The four components shown separately for clarity. (C) Example component time traces, color coded to match panel B. Local temporal averages for three time-points, each with the components displayed with intensity weighted by the respective integrated fluorescence over that time-window, and the local video average of the same time window show that the full axonal group is active when the components have high activity levels. (D) Suite2p results on the same data breaks up the axons into multiple segments, similar to dendritic imaging [17].

## 7 Conclusions and Discussions

In this work we build on the GraFT methodology in several ways. First, we introduce computational improvements that increase the speed of component identification.

Second, we improve the usability of GraFT through a compiled graphical user interface.

Finally, we demonstrate the extent of GraFT’s ability to identify components in novel large-scale imaging applications, including vascular and axonal imaging.

The computational advances we present target the bottle necks of performing the weighted *ℓ*_1_ regularized optimization in the spatial profile estimation and the least-squares time trace update. For the former, we tested multiple solvers, identifying significant gains by switching to an MPC solver over a more traditional IPM. For the latter, we implemented a compressive computation, reducing the effective size of the data in the least squares problem and therefore the computation time.

The comparative analysis of ASM and IPM solvers for the LASSO optimization within the GraFT algorithm reveals distinct tradeoffs that are highly dependent on dataset characteristics. The ASM solver, implemented via MATLAB’s mpcActiveSetSolver, consistently demonstrated faster runtimes compared to the IPM-based quadprog approach, particularly for small to medium-sized data windows. This improved performance is largely attributed to its effective warm-start strategy and lower per-iteration computational cost. While IPM methods benefit from robust convergence properties and are theoretically well-suited for large-scale problems due to fewer iterations, their higher per-iteration computational expense becomes a limiting factor when applied to smaller or less complex datasets. In contrast, the ASM method’s ability to rapidly adjust its working set of constraints makes it a more efficient choice in scenarios where rapid, repeated optimizations are required.

While the speedups afforded by changing the LASSO optimization are direct replacements of specific steps in the code, the compressive computation changes the data size and magnitudes, therefore changing the balance of sparsity and data variance. We thus recommend tuning the sparsity coefficient, *λ*, and normalized compressed GraFT in a majority of cases. Normalizing GraFT in this way provides much better stability at higher compression levels and spatial profile comparison, making it the appropriate choice for larger data that can take the most advantage of high compression levels. However, in a specific case where time traces information is strongly prioritized over spatial information, non-normalized GraFT remains a good alternative due to its strong self-consistency of trace preservation up to 64X compression and its ability to identify a more comprehensive collection of time traces.

Our newly developed GUI significantly enhances the accessibility and efficiency of GraFT by providing user-friendly parameter configuration, algorithm execution, and preliminary result filtering within a single application. Both the stand-alone and MATLAB-based versions are also helpful in maintaining flexibility with existing pipelines. Future releases will expand data-format support, including Neurodata Without Borders (NWB) compatibility [40], to further extend the platform’s utility and maintain its user-oriented design.

## Acknowledgments

This work was supported by funding from the National Institutes of Health (R01 EB026936 to GM, R01 DC018650 to KVK, U19 NS123717 to AEB, TB, DK and GM, R35 NS097265 to TB and DK), the National Science Foundation (CAREER 2145247 to KVK), a Mathworks MATLAB Community Toolbox Training fellowship to AEB, a KAVLI NDI distinguished postdoctoral fellowship to JL and an Early (P2SKP3 164948) and an Advanced (P300PA 177804) Postdoctoral Mobility fellowship from the Swiss National Science Foundation to JB.

The authors would like to thank Jacob Duckworth for his assistance with the vasculature dataset.

## A Data

### NeuroFinder

We analyzed one of the NeuroFinder datasets [36], which contained two-photon calcium imaging at the somatic level, as well as manually annotated neuron spatial profiles. We ran GraFT on a 445*µ*m × 445*µ*m field of view in area vS1 of mouse visual cortex, recorded at 8 Hz.

### NAOMi

For evaluation of compression on GraFT, we generated both the (490 × 490 × 100) *µ*m × 9,000 time point (equivalent to 3 minutes) and (500 × 500 × 100) *µ*m× 20,000 time point (approximately equivalent to 11 minutes) in NAOMi with a sampling rate of 30Hz (`spike_opts.dt = 1/30`). Specifically, the imaging was simulated using a 0.6NA Gaussian beam (`psf_params.NA = 0.6`), 0.8NA objective (`psf_params.objNA = 0.8`), and 40 mW power (`tpm_params.pavg = 40`). The imaging depth was set to the middle of the simulated volume (`vol_params.vol_depth = 100`).

### Vasculature

We analyze a single trial from the dataset collected in [38]. Imaging was performed in mice with GCaMP8.1 in arteriole smooth muscles at 4.47 Hz. We used an extended thinned skull window that spanned an approximately 10 mm by 10 mm region of skull over the cortical mantle. To create the pial arteriole mask, we took the standard deviation over time of the trial’s first 1250 diffusion-filtered GCaMP8.1 fluorescence images. The result was normalized and contrast-enhanced using MATLAB’s “adapthisteq” function. Adobe Photoshop was used to sharpen, threshold, and manually connect pial arteriole segments that fell below the threshold. The thin-skull window edge was then manually defined and saved as the “rim” mask. The pial arteriole mask was used to define a vessel graph using procedures developed for reconstruction of the 3-dimensional microvascular connectome [41]. Briefly, the two-dimensional mask was skeletonized and diameter was estimated using MATLAB’s “bwskel” and “bwdist” respectively. Skeleton pixels, which define the vessel centerline, were classified as either an endpoint, a link that connects two neighbors, or a node that connects three or more neighbors. Skeleton pixels were downsampled and dilated by the vessel diameter to define skeleton segments. We then calculated the raw fluorescence signal *F* (*x, t*) as the median intensity of pixels within each skeleton segment at each time point. Finally, 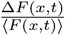 was calculated and stored for subsequent analyses, where Δ*F* (*x, t*) = *F* (*x, t*) *−* ⟨*F* (*x, t*) ⟩ and ⟨*F* (*x, t*) ⟩ denotes the time-averaged intensity at each location *x*. Additional experimental details can be found in [38].

### Axonal data

Cholinergic axonal imaging data were collected as described in [39] from the auditory cortex of ChAT-Cre mice (stock no. 006410, Jackson Laboratory; 2-3 months old). To label cholinergic neurons and their axonal projections, an AAV vector encoding the calcium indicator Axon-GCaMP6s (1 *µ*L;

AAV5-hSynapsin1-FLEx-axon-GCaMP6s, Addgene) was injected into the left basal forebrain (coordinates from bregma: AP -0.5 mm, ML 1.8 mm, DV 4.5 mm). To express a calcium indicator in auditory cortical neurons, an AAV vector encoding jRGECO1a (∼ 0.8–1.5 *µ*L; AAV1-syn-jRGECO1a, Addgene) was injected into layer 2/3 of the left auditory cortex. A 3 mm circular glass window (Warner Instruments) was implanted over the exposed cortical surface to permit optical access (centered 1.75 mm anterior to lambda on the ridge line). Two-color, two-photon imaging of cholinergic axons and auditory cortical neurons was performed using a resonant-scanning two-photon microscope (Neurolabware) equipped with a X16 objective (Nikon), angled at 45–55^*?*^, and a Spectra-Physics Insight X3 laser (980 nm excitation, ≤ 40 mW laser power). A tunable lens enabled near-simultaneous imaging of two planes—layer 1 (60–100 *µ*m below dura) and layer 2/3 (150–200 *µ*m below dura)—with a field of view of 312 × 192 *µ*m^2^ per plane at 15.63 Hz. Animals underwent longitudinal two-photon imaging daily during learning of an auditory discrimination task (see [42, 43]). The data analyzed in this manuscript correspond to the layer 1 plane from the first day of learning in one animal (male, 2 months old at imaging start), comprising 40,807 frames from the green channel (cholinergic axons only).

## B Implementation

### B.1 ethod hyper-parameters

The following GraFT hyper-parameters were kept consistent across all datasets. We enforced nonnegativity in both the dictionary and coefficients (`nneg_dict = true and nonneg = true`), and used `grad_type = ‘full ls cor’` with `maxiter = 0.01, tolerance = 1e-8`, and `likely_form = ‘gaussian’` to perform gradient-based optimization. The gradient step-size parameters were `step_s = 1` and `step_decay = 0.995`, with a maximum of `max_learn = 50` learning iterations and a convergence tolerance of learn eps = 5e-4. Each update used GD iters = 1 gradient-descent iteration. We disabled embedding updates (`updateEmbed = false`), but maintained spatial normalization (`normalizeSpatial = true`). Lastly, we reduced dimensionality in the correlation kernel (reduce dim = true) and used a graph embedding approach (`corrType = ‘embedding’`).

#### NeuroFinder

For the NeuroFinder dataset, the primary regularization parameters were set to *λ* = 0.7, *λ*_Forb_ = 0.2, and *λ*_Corr_ = 0.1. Spatial patching (`patchGraFT = true`) was enabled only during specific runtime tests (where the size of the dictionary was varied). All other parameters were retained hyper-paramters listed above.

#### Vasculature

For the vasculature dataset, the same baseline hyper-parameters were used, but the regularization terms were updated to *λ*_Forb_ = 0, *λ*_Cont_ = 1, and *λ*_Corr_ = 0.5. All other settings, including the correlation kernel, remained unchanged.

#### Axonal data

For the axonal data we used the parameters *λ* = 2, *λ*_Forb_ = 0.2, *λ*_Cont_ = 0.1, and *λ*_Corr_ = 0.1. All other settings, including the correlation kernel, remained unchanged.

### B.2 Computing Environment

The solver optimization analyses were done on a Windows PC tower with 13th Gen Intel(R) Core(TM) i7-13700K 3.4 GHz processor, 64 GB of DDR4 RAM and a GeForce3080 GPU. We wrote custom MATLAB scripts to iteratively test and time each GraFT.m and patchGraFT.m scripts with the different solvers.

### B.3 Solver comparison in optimization

To improve computational performance, we implemented two separate quadratic programming solvers tailored to the structure of our LASSO subproblem: MATLAB’s Optimization Toolbox `quadprog` solver, and the MPC Toolbox’s `mpcActiveSetSolver`. The former is a general-purpose convex QP solver that supports various algorithms, including interior-point and active-set methods, and is widely used across a range of convex problems. In contrast, `mpcActiveSetSolver` leverages the KWIK algorithm, a reduced-Hessian active-set method originally developed for real-time control applications. The KWIK method, based on the dual Goldfarb-Idnani algorithm and optimized for reduced Hessian SQP [34], offers computational advantages in problems with few degrees of freedom but many constraints—conditions often present in our L1-constrained formulation. While `quadprog` ensures generality and numerical stability, the specialized structure and warm-start capabilities of `mpcActiveSetSolver` offer a substantial speedup, particularly in iterative regimes where active sets change slowly.

We exploit both solvers depending on problem size and conditioning, with `mpcActiveSetSolver` providing the greatest gains in large-scale regimes with structured sparsity.

**Fig S1.**
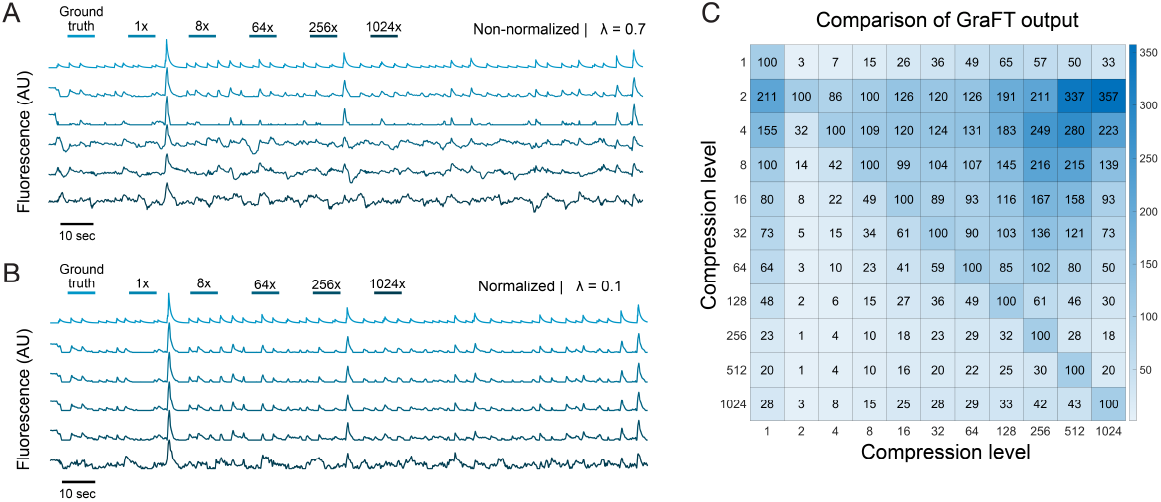
Additional influences of GraFT compression level on identified time trace components. Example medium correlated time traces (0.4 *< ρ <* 0.7) across compression levels in (A) non-normalized GraFT and (B) normalized GraFT. (C) Heatmap for normalized GraFT results across compression levels. Values in the heatmap are the actual number of identified traces.

## Footnotes

1 Users are encouraged to consult the GitHub GraFT wiki site for detailed tutorials on how to employ preprocessing features, parameter selection, and analysis of GraFT results.

